# Proteomic profiling of whole tissue sections in cardiac ATTR amyloidosis reveals increased extracellular matrix remodeling

**DOI:** 10.64898/2026.04.01.715792

**Authors:** Annelore Vandendriessche, Teresa M Maia, Frank Timmermans, Delphi Van Haver, Sara Dufour, An Staes, Joost Schymkowitz, Frederic Rousseau, Rodrigo Gallardo, Michel Delforge, Jo Van Dorpe, Simon Devos, Francis Impens, Amélie Dendooven

## Abstract

Cardiac transthyretin amyloidosis (ATTR-CA) is caused by myocardial deposition of misfolded transthyretin, leading to progressive heart failure. Disease pathology, however, extends beyond passive amyloid deposition and also involves active processes such as extracellular matrix (ECM) remodeling and immune activation.

Mass spectrometry (MS) is the gold standard for amyloid typing in diagnostics. Here, we applied quantitative MS-driven proteomics on formalin-fixed paraffin-embedded whole cardiac tissue sections from six ATTR-CA cases, ten unaffected controls and four AL-CA controls to investigate protein expression changes. In addition to transthyretin, over 500 proteins were upregulated in ATTR-CA biopsies, including complement and coagulation factors as well as extracellular matrix (ECM) remodeling proteins.

Among these, members of the A Disintegrin and Metalloproteinase with Thrombospondin Motifs (ADAMTS) family, metalloproteinases (MMPs), and Tissue Inhibitor of Metalloproteinases (TIMP3) showed significant upregulation. These proteins are key regulators of ECM turnover and structural integrity. Immunohistochemistry confirmed ADAMTS4 enrichment in amyloid deposits, while TIMP3 showed strong expression in cardiomyocytes and weaker staining within amyloid deposits.

Together, these findings indicate that ECM remodeling, alongside complement and coagulation activation, represents a reproducible feature of cardiac ATTR amyloidosis. Whole-tissue proteomics provides biological insights that extend beyond amyloid typing, with potential implications for biomarker discovery and therapeutic targeting in ATTR-CA.

## INTRODUCTION

Amyloidosis is a group of rare diseases characterized by extracellular deposition of misfolded proteins forming fibrils. Transthyretin amyloidosis (ATTR) is one of the most prevalent disease subtypes, out of the approximately 42 that have been identified to date ^1^. Under physiological conditions, TTR circulates in the bloodstream as a stable tetramer, transporting thyroxine and vitamin A. During amyloidogenesis, however, the tetramer dissociates into unstable monomers that misfold and aggregate into insoluble fibrils which progressively accumulate in tissues, mainly the heart. ATTR exists in two forms: wild-type ATTR (ATTRwt) and hereditary ATTR variants (ATTRv) ^2,3^. Both forms result in significant morbidity and mortality from progressive heart failure with preserved ejection fraction (HFpEF) due to increased myocardial stiffness and impaired ventricular relaxation ^4^. Studies have shown that ATTRwt contributes to diastolic dysfunction and is present in up to 13% of HFpEF cases in the elderly ^5^. On the cellular level, TTR fibrils and soluble aggregate forms can exert direct cytotoxic effects, inducing oxidative stress, dysregulation of calcium handling, and apoptosis in cardiomyocytes ^6^. In parallel, remodeling of the extracellular matrix (ECM) compromises myocardial elasticity through increased deposition and turnover of matrix components ^7^.

Key regulators of ECM homeostasis and organization are metallopeptidases and their inhibitors. These include Matrix Metalloproteinases (MMP), members of the A Disintegrin and Metalloproteinase (ADAM) and of the ADAM with Thrombospondin Motifs (ADAMTS) family, and the Tissue Inhibitors of Metalloproteinases (TIMP). MMPs are known for their ability to degrade ECM components, making them important regulators of myocardial structure and function ^8^. Amyloid fibril deposition may disrupt ECM dynamics, as increased expression of several MMPs (MMP-2 and -9, more specifically) and TIMPs has been reported in cardiac light chain amyloidosis (AL-CA) but not in ATTR-CA ^9^. Beside MMPs, ADAMs and ADAMTSs are also important proteases in cardiovascular ECM homeostasis. Specifically, they modulate ECM composition by regulating proteoglycan turnover ^10^. In heart failure, ECM remodeling involves not only collagen turnover but also altered proteoglycan metabolism. This leads to proteoglycan accumulation, which contributes to myocardial stiffness and dysfunction ^11^.

The activity of metallopeptidases is regulated by their endogenous TIMP inhibitors. The balance between the activity of MMPs, ADAMs, ADAMTSs and TIMPs plays a role in the stability and normal function of the ECM. Among these, TIMP3 is unique for its strong binding to the ECM and broad inhibitory profile, making it a key protector of myocardial ECM integrity ^12^. TIMP3 selectively inhibits several ADAMTS family members, specifically ADAMTS1, -2, -4 and -5 ^13^.

Contemporary mass spectrometry (MS)-based proteomics allows for accurate amyloid typing and has been used in specialized centers worldwide ^14^. In their most advanced form, these analyses comprise laser capture microdissection (LCM) of amyloid plaques from patient tissue sections, followed by liquid chromatography-tandem mass spectrometry (LC-MS/MS) analysis ^15^. However, MS-based analysis of whole tissue sections, without LCM, containing also cells and ECM surrounding the amyloid plaques, is suitable to investigate changes in the amyloid-associated proteome ^16,17^.

In the present study we analyzed whole cardiac biopsies from ATTR-CA patients and unaffected controls by LC-MS/MS and analyzed the data with particular attention to ECM components including metalloproteinases and their inhibitors. Along with complement and coagulation factors, we observed significantly increased expression of various ECM remodeling factors. We confirmed the upregulation of ADAMTS4 and TIMP3 by immunohistochemistry (IHC), both key regulators of matrix turnover and stability, highlighting the important role of ECM remodeling in cardiac amyloidogenesis.

## MATERIALS AND METHODS

### Sample collection and ethics approval

After obtaining approval from the Ghent University Hospital’s Ethics Committee (EC: ONZ2022-0301), 6 cardiac formalin-fixed paraffin-embedded (FFPE) ATTR biopsies (cases A1-A6) and 10 unaffected control biopsies (cases C1-C10) were selected. The ATTR cohort consisted of male patients (age range 41-78 years). Nine subjects of the control group were male, one female (age range 42-73 years). The youngest patient in the ATTR group was known with familial amyloid polyneuropathy. For two other patients, clinical diagnosis was of wild-type ATTR without genetic testing, and in three additional cases, no pathogenic mutation in the TTR gene was detected (**Table 1**). For comparative analyses, four cardiac AL (light chain) amyloidosis biopsies (three AL lambda and one AL kappa) were included.

**Table 1.** Demographic and clinical characteristics of included case and control biopsies.

| Identifier | Age | Gender | Histology diagnosis | CR staining | TTR staining | Biopsy indication | Year of biopsy | TTR mutation status |
| --- | --- | --- | --- | --- | --- | --- | --- | --- |
| A1 | 64 | M | ATTR | + | + | Amyloidosis suspected | 2014 | Wild-type |
| A2 | 41 | M | ATTR | NC* | + | Amyloidosis suspected | 2015 | Ala59Val (heterozygous) |
| A3 | 75 | M | ATTR | + | + | Amyloidosis suspected | 2015 | Wild-type |
| A4 | 68 | M | ATTR | + | + | Amyloidosis suspected | 2018 | Not tested |
| A5 | 78 | M | ATTR | + | + | Amyloidosis suspected | 2021 | Wild-type |
| A6 | 74 | M | ATTR | + | + | Amyloidosis suspected | 2022 | Not tested |
| C1 | 43 | M | Normal (0R) | - | N/A** | Post HTX*** | 2015 | N/A |
| C2 | 53 | M | Normal (0R) | - | N/A | Post HTX | 2015 | N/A |
| C3 | 52 | M | Normal | - | N/A | Congestive CM suspected (excluded) | 2015 | N/A |
| C4 | 42 | F | Normal (0R) | - | N/A | 3rd post HTX | 2016 | N/A |
| C5 | 62 | M | Normal (0R) | - | N/A | Post HTX | 2017 | N/A |
| C6 | 51 | M | Normal (0R) | - | N/A | 4m post HTX | 2019 | N/A |
| C7 | 45 | M | Normal (0R) | - | N/A | 2,5m post HTX | 2019 | N/A |
| C8 | 73 | M | Normal (0R) | - | N/A | Post op HTX | 2021 | N/A |
| C9 | 69 | M | Normal | - | - | Amyloidosis suspected (excluded) | 2022 | N/A |
| C10 | 65 | M | Normal (0R) | - | N/A | First control after combined HTX/KTX | 2022 | N/A |
| AL1 | 55 | M | AL lambda | + | - | Amyloidosis suspected | 2014 | N/A |
| AL2 | 75 | M | AL lambda | + | - | Amyloidosis suspected | 2017 | N/A |
| AL3 | 66 | M | AL lambda | + | - | Amyloidosis suspected | 2021 | N/A |
| AL4 | 49 | F | AL kappa | + | - | Amyloidosis suspected | 2014 | N/A |

\*NC: inconclusive, \*\*N/A: not applicable, \*\*\*HTX: heart transplant.

### Histology and immunohistochemical analysis

Biopsy slides were stained for H&E, Congo Red (CR), and immunohistochemistry (IHC) and examined by two pathologists (AV and AD). ATTR cases were confirmed by CR and TTR IHC (BSB3621, BioSB, Santa Barbara, CA, USA). The control cases consisted of histologically normal post-transplant biopsies (n=8, all graded 0R (18)), histologically normal native myocardial biopsies (n=2), and cardiac AL amyloidosis biopsies (n=4).

Immunohistochemistry for ADAMTS4 and TIMP3 was performed on a control biopsy and on cardiac tissue from the explanted heart of case A3 from the ATTR-CA cohort. This was done on 3 μm FFPE sections using the BenchMark Ultra automatic immunostainer and the ultraView Universal DAB Detection Kit (both Ventana Medical Systems, Roche Diagnostics, Basel, Switzerland). For ADAMTS4, a mouse monoclonal antibody (OTI6F8, MA5-26715, Thermo Fisher Scientific, Waltham, MA, USA, dilution 1:400) was used. Heat-induced epitope retrieval was performed with Cell Conditioning 1 (95 °C, 8 min). Sections were baked and deparaffinized, after which the primary antibody was applied manually.

Counterstaining was carried out with Hematoxylin II and Bluing reagent. For TIMP3, a rabbit polyclonal antibody (ab93637, Abcam, Cambridge, UK, dilution 1:400) was used without antigen retrieval. Subsequent processing, including section preparation and counterstaining, was performed as described for ADAMTS4.

### Proteomics sample preparation

For each case, a 10 µm thick tissue section was collected using a microtome (HM340E, Thermo Fisher Scientific, Waltham, MA, USA) in an Eppendorf tube (AdhesiveCap 500 opaque, Zeiss, Oberkochen, Germany), with tissue areas ranging from 2,42 to 11,98 mm^2^. For the ATTR group, the amyloid load (defined as the proportion of tissue area occupied by Congo red-positive deposits) ranged from 21 to 83.5% (**Table 2**). Samples were prepared using an in-house protocol ^18^, adapted from the HYPERsol method ^19^. Proteins were extracted by adding 5% sodium docedyl sulfate (SDS) in 50 mM triethylammonium bicarbonate buffer (TEAB) and incubating overnight at 50°C. This was followed by ultrasonication and incubation at 80°C to break formaldehyde crosslinks. Cysteine bridges were broken through reduction and alkylation with 15 mM dithiothreitol (DTT) and 30 mM iodoacetamide (IAA), respectively. Phosphoric acid was added to a final concentration of 1.2 % and samples were diluted 7-fold with binding buffer containing 90 % methanol in 100 mM TEAB, pH 7.55.

**Table 2.** Sample size, amyloid load, and cellularity of ATTR biopsy specimens.

| Identifier | Sample size (mm<sup>2</sup>) | Absolute amyloid load (mm<sup>2</sup>) | Relative amyloid load (%) | Total number of nuclei | Number of nuclei per mm<sup>2</sup> area |
| --- | --- | --- | --- | --- | --- |
| A1 | 5,71 | 4,77 | 83,5 | 6626 | 1160,4 |
| A2 | 2,42 | 1,111 | 45,9 | 2066 | 853,7 |
| A3 | 11,979 | 8,984 | 75 | 13397 | 1118,4 |
| A4 | 5,43 | 1,141 | 21 | 4231 | 779,2 |
| A5 | 5,121 | 1,987 | 38,8 | 3582 | 699,5 |
| A6 | 8,469 | 2,524 | 29,8 | 11231 | 1326,1 |

Proteins were trapped on S-trap columns, washed with binding buffer, 50 % methanol in chloroform and digested on-column with 1 μg trypsin at a 0.05 µg/µl concentration by incubation at 37°C for 16 hours. After incubation, peptides were eluted from the S-trap column with 3 elution buffers, first with 80 µL 50 mM TEAB, then with 80 μL 0.2% formic acid (FA) in water and finally with 80 µL 0.2 % FA in water/acetonitrile (ACN) (50/50, v/v). Eluted peptides were dried completely by vacuum centrifugation, redissolved in 0.1 % TFA in water/ACN (98/2, v/v) and 1/10^th^ of the peptide solution was loaded on Evotips (Evotip Pure, Evosep, Odense, Denmark) for LC-MS/MS analysis.

### LC-MS/MS and data analysis

Samples were run in data-independent acquisition parallel accumulation serial fragmentation (DIA-PASEF) mode using an Evosep One LC system (Evosep, Odense, Denmark) in-line connected to a TimsTOF SCP (Bruker, Billerica, MA, USA). Peptides were analyzed with the 20 SPD whisper method using the Aurora Gen3 Elite column (15 cm x 75 µm I.D., 1.7 µm beads, IonOpticks, Fitzroy, VIC, Australia), heated to 45°C. Peptides were eluted from the column through the predefined 20 SPD whisper gradient, using 0.1% formic acid (FA) in LC-MS-grade water as solvent A and 0.1% FA in acetonitrile (ACN) as solvent B. Eluting peptides were measured in positive polarity with a full-scan range of 100 m/z to 1700 m/z. The trapped ion mobility spectrometry (TIMS) was operated at a fixed duty cycle close to 100%, a ramp and accumulation time of 100 ms, ranging from 1/K0 = 0.64 Vscm² to 1/K0 = 1.5 Vscm². Collision energy was linearly ramped as a function of the inverse mobility from 20 eV at 1/K0 = 0.60 Vscm² to 59 eV at 1/K0 = 1.60 Vscm². A DIA-PASEF mass range of 400 Da to 1000 Da was used in a mobility range of 1/K0 = 0.64 Vscm² to 1/K0 = 1.37 Vscm² using a window size of 25 Da, resulting in a cycle time of 0.96 s (**Table 3**).

**Table 3.** DIA-PASEF windows.

| Cycle Id | Start IM [1/K0] | End IM [1/K0] | Start Mass [m/z] | End Mass [m/z] |
| --- | --- | --- | --- | --- |
| 1 | 0.64 | 0.83 | 400 | 425 |
| 2 | 0.64 | 0.85 | 425 | 450 |
| 3 | 0.64 | 0.87 | 450 | 475 |
| 4 | 0.64 | 0.9 | 475 | 500 |
| 5 | 0.64 | 0.92 | 500 | 525 |
| 6 | 0.64 | 0.94 | 525 | 550 |
| 7 | 0.64 | 0.97 | 550 | 575 |
| 8 | 0.64 | 0.99 | 575 | 600 |
| 1 | 0.83 | 1.01 | 600 | 625 |
| 2 | 0.85 | 1.04 | 625 | 650 |
| 3 | 0.87 | 1.06 | 650 | 675 |
| 4 | 0.9 | 1.09 | 675 | 700 |
| 5 | 0.92 | 1.11 | 700 | 725 |
| 6 | 0.94 | 1.13 | 725 | 750 |
| 7 | 0.97 | 1.16 | 750 | 775 |
| 8 | 0.99 | 1.18 | 775 | 800 |
| 1 | 1.01 | 1.37 | 800 | 825 |
| 2 | 1.04 | 1.37 | 825 | 850 |
| 3 | 1.06 | 1.37 | 850 | 875 |
| 4 | 1.09 | 1.37 | 875 | 900 |
| 5 | 1.11 | 1.37 | 900 | 925 |
| 6 | 1.13 | 1.37 | 925 | 950 |
| 7 | 1.16 | 1.37 | 950 | 975 |
| 8 | 1.18 | 1.37 | 975 | 1000 |

LC-MS/MS runs of all samples were searched using the DIA-NN algorithm (version 1.8.1) ^20^, with primarily default settings, including a false discovery rate set at 1% on precursor and protein levels. Spectra were searched against the human canonical reference proteome sequences (downloaded from www.uniprot.org, database release version of 2024_01), comprising 20,596 sequences, and common contaminants.

Trypsin was selected as digestion enzyme. Variable modifications included carbamidomethylation of cysteines, oxidation of methionine, and acetylation of N-termini, with a limit of five variable modifications per peptide. The precursor mass range was restricted to 400-1000 m/z, and the match between runs (MBS) feature was activated. The mass accuracy was set to 20 ppm for MS2 and 15 ppm for MS1.

For statistics, an in-house script in R programming language (version 4.1.1) was used. The main output table from DIA-NN was filtered with a precursor and protein library q-value threshold of 1%, retaining only proteins with at least one proteotypic peptide. Peptide and protein ID rates were based on features detected in at least 2 samples. For differential abundance analysis, LFQ-normalized protein intensities were log2 transformed. Proteins with fewer than 3 valid data points in any experimental condition were excluded. Missing data were imputed by random sampling from a normal distribution centered around each sample’s noise level, using the package DEP ^21^. Pairwise comparisons between the ATTR and unaffected group were performed using the limma package ^22^, with statistical significance for differential regulation set at an FDR cut-off value of 0.05 and a |log2(fold change)| >= 2.

Proteins that were completely absent in one of both conditions were added as regulated hits for unsupervised clustering and heatmap visualization. For relative abundance analysis per sample and ranking of TTR and other amyloidogenic and signature proteins, iBAQ values were calculated using the get_iBAQ function from DIAgui package.

### Gene ontology enrichment and network analysis

Gene ontology enrichment analysis was performed with the DAVID online tool ^23^, using the significantly up- or downregulated proteins as input. Filtering for GO terms was carried out with an FDR for the enrichment test <0.05 (**Supplementary Table S4**).

### Additional data analysis

Quality control analyses were performed to assess data integrity and sample comparability. Main feature intensity distributions were visualized as boxplots using the limma package to identify potential outliers or differences in overall signal intensity. A Venn diagram was constructed using the Bioinformatics & Evolutionary Genomics online platform from VIB/Ghent University (https://bioinformatics.psb.ugent.be), allowing visualization of protein identifications between sample groups (**Supplementary Table S5**). In addition, sample-to-sample correlations were calculated based on protein expression values and visualized as a heatmap after hierarchical clustering. Plotted values represent Spearman correlation coefficients between all sample pairs. The mass spectrometry proteomics data have been deposited to the ProteomeXchange Consortium via the PRIDE partner repository with the dataset identifier PXD072972 ^24^.

## RESULTS

### Quantitative proteome analysis of ATTR-CA samples

To measure protein changes associated with ATTR amyloidosis in the heart, we performed an in-depth proteome analysis on myocardial biopsies from six ATTR-CA amyloidosis patients, 10 unaffected control subjects and 4 AL controls. Histological examination confirmed the presence of amyloid deposits in ATTR cases by hematoxylin and eosin, Congo red, and TTR IHC (**Figure 1A**). As a next step, 10 µm thick tissue sections of each FFPE biopsy were subjected to MS-based proteome analysis. Proteins from each section were extracted using a protocol based on SDS solubilization in combination with ultrasonication ^18^. Extracted proteins were digested with trypsin and the resulting peptide mixture was analyzed by label-free LC-MS/MS analysis (**Figure 1B**). In total, 4,291 proteins were identified (**Supplementary Table S1, Supplementary Table S6**), of which 3,824 were reliably quantified with protein intensity values in at least three replicates of at least one condition (**Supplementary Table S2**). Statistical analysis and differential expression analysis revealed 519 upregulated and 75 downregulated proteins in the ATTR samples compared to the unaffected controls. These proteins were visualized in a heatmap after unsupervised hierarchical clustering (**Figure 1C**, **Supplementary Table S3**).

**Figure 1.**
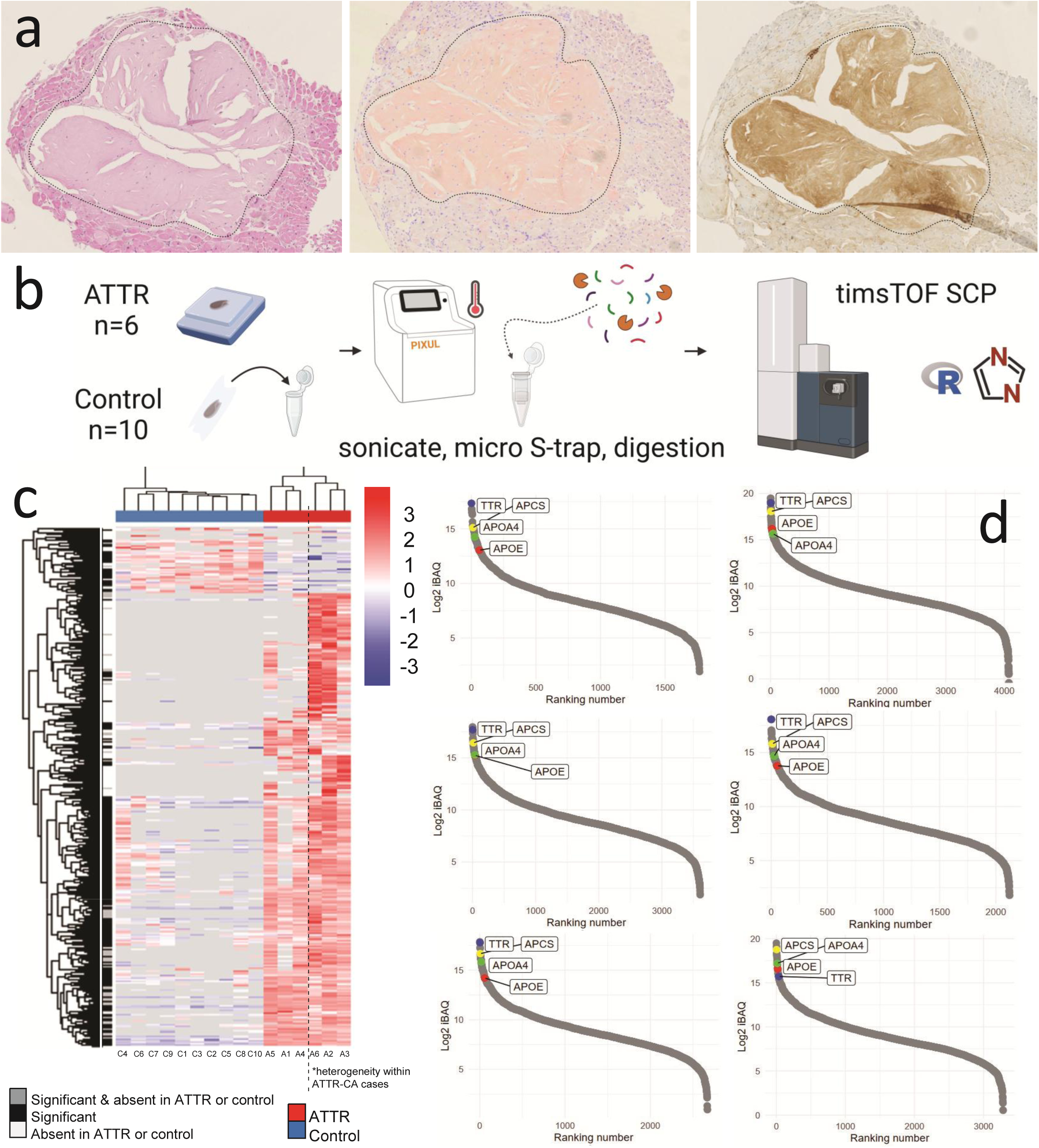
(A) Myocardium with ATTR deposits (Case A6). FFPE (formalin-fixed paraffin embedded) tissue stained with Haematoxylin and Eosin (left), Congo red (middle) and TTR IHC (right). The deposition is indicated by a dotted line. Original magnification 2.5x. **(B) Schematic representation of the workflow.** Case selection, preparation of FFPE tissue sections, sample processing according to laboratory protocol, LC-MS/MS analysis, and data analysis with DIA-NN and R (left to right). **(C) Heatmap of differentially expressed proteins after unsupervised hierarchical clustering.** Control cases are shown on the left and ATTR cases on the right. The top cluster represents 75 proteins that were downregulated in the ATTR cohort, whereas the bottom cluster comprises the 519 upregulated proteins. Within the ATTR group, the dotted line highlights a subgroup on the right in which additional protein upregulation was observed in some, but not all, cases. **(D) ATTR protein abundance curves.** From each of the six ATTR samples, scatter plots show protein intensities ranked by relative abundance. Protein intensities of TTR (blue) and amyloid signature proteins APCS (yellow), ApoE (red) and ApoAIV (green) are highlighted.

### Upregulation of TTR and amyloid signature proteins

In the ATTR group, TTR showed the highest relative abundance of all quantified amyloidogenic proteins. Indeed, when visualized in a protein abundance ‘S-curve’ plot, TTR is positioned among the top-ranked proteins for every ATTR sample (**Figure 1D**). Besides TTR, we observed significant upregulation of other amyloidogenic proteins including lysozyme C, apolipoprotein CIII, apolipoprotein AII, and apolipoprotein AIV (ApoAIV).

ApoAIV is not only an amyloidogenic precursor but also an amyloid signature protein. Other known fibril-forming proteins associated with systemic forms of amyloidosis, including immunoglobulin light chain lambda and kappa, serum amyloid A (SAA), β2 microglobulin, apolipoprotein AI, gelsolin and fibrinogen alpha, were present in both ATTR and healthy samples without significant difference (**Supplementary Table S2**). Moreover, amyloid signature proteins apolipoprotein E (ApoE), ApoAIV and serum amyloid P component (SAP) showed high intensities in the ATTR samples compared to the controls and were detected in all ATTR samples (**Figure 1D**). Vitronectin and clusterin, proteins well known to be associated with amyloidosis, also showed significant upregulation.

### Functional annotation and pathway analysis of amyloid-associated proteins

A gene ontology (GO) analysis was performed to gain insight into functional pathways enriched in ATTR-CA, using the significantly upregulated proteins as input. This revealed enrichment of biological process terms related to immune and complement activation, while cellular component and molecular function terms indicated remodeling of the ECM (**Figure 2A**). To refine the interpretation, complement proteins, coagulation factors and ECM remodeling proteins, including members of the ADAMTS family were manually selected from the list of significantly regulated proteins and their intensities were visualized in separate heatmaps after unsupervised clustering (**Figure 2B-D**).

**Figure 2.**
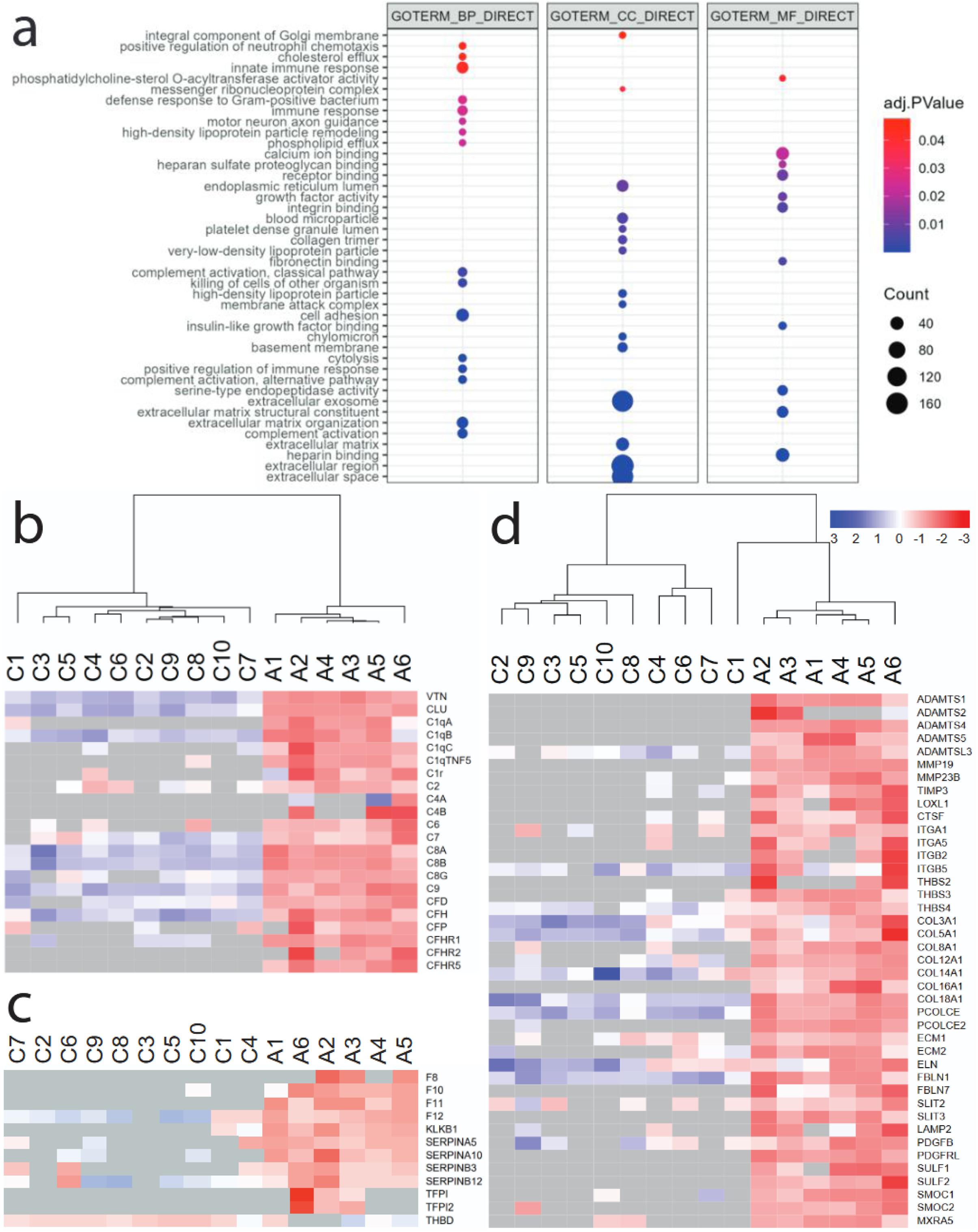
(A) Gene ontology (GO) analysis. Pathways enriched in ATTR-CA are related to immune and complement activation, and remodeling of the ECM. The 519 significantly upregulated proteins were used as input. **(B-D) Heatmaps of selected proteins.** Heatmaps illustrating differential expression of proteins associated with the complement system (B), coagulation cascade (C), and ECM remodeling (D). In each heatmap, controls are shown on the left and ATTR cases on the right.

### Enrichment of complement and coagulation factors

C1q subunit A (C1qA), B (C1qB) and C (C1qC) were significantly upregulated in ATTR samples. Additional complement components were also identified (**Supplementary Table S1**), of which C2, C4A and C4B, C6, C7, C8 alpha, beta, and gamma chain, C9, C1R, complement factor D (CFD) and H (CFH), and complement factor H-related proteins (CFHR) 1, 2, 3 and 5 were significantly upregulated (**Figure 2B**). Furthermore, we identified scavenger receptor c-type lectin (SRCL) collectin-12 in two ATTR samples, and in none of the control tissues, however, collectin-12 itself could not be reliably quantified.

Additionally, several components of the coagulation pathway were significantly upregulated in the amyloidosis group, including coagulation factor X (F10) of the common pathway, and VIII (F8), XI (F11), and XII (F12) of the intrinsic pathway. Also, kallikrein B1 (KLKB1), tissue factor pathway inhibitor (TFPI) 1 and 2, tissue-type plasminogen activator (PLAT), plasma serine protease inhibitor (SERPINA5), protein Z-dependent protease inhibitor (SERPINA10), serpin B3 (SERPINB3) and serpin B12 (SERPINB12) were upregulated, while thrombomodulin (THBD) was downregulated in ATTR samples (**Figure 2C**).

To check whether these amyloid-associated changes are specific to ATTR-CA, we additionally analyzed four AL-CA samples derived from three AL lambda and one AL kappa patient. Interestingly, enrichment of complement and coagulation pathways was also present in AL, but the magnitude of upregulation was generally less pronounced compared to ATTR, indicating that the amyloid-associated proteome might differ between amyloid types and that the enrichment of these factors is specific to ATTR (**Supplementary Figure 1, Supplementary Table S7**).

### Enrichment of ECM remodeling factors

ADAMTS1, ADAMTS4, ADAMTS5, and ADAMTS-like protein 3 (ADAMTSL3) showed a consistent and significant upregulation in ATTR samples, while from the ADAM family no proteins were differentially expressed. In addition, MMP19 and MMP23B were upregulated, along with TIMP3 and thrombospondin 2, 3 and 4 (THBS2, 3, 4), as components of the ECM that interact with MMPs. We further observed increased presence of several collagens (COL3A1, COL5A1, COL6A1, COL8A1, COL12A1, COL14A1, COL16A1, and COL18A1) as well as procollagen C-endopeptidase enhancer (PCOLCE) and PCOLCE2. Finally, extracellular matrix protein (ECM) 1 and 2 were upregulated together with other factors contributing to ECM remodeling, including cathepsin F (CTSF), lysosome-associated membrane glycoprotein (LAMP) 2, and matrix-remodeling associated protein 5 (MXRA5) (**Figure 2D**). Similar as for the complement and coagulation factors, extended analyses including AL cases showed that the upregulation of ECM remodeling factors was most pronounced in ATTR (**Supplementary Figure 1, Supplementary Table S7**).

### Validation of increased ADAMTS4 and TIMP3 expression

To confirm the MS-based detection of ECM remodeling factors we selected ADAMTS4 and TIMP3 as targets for antibody-based detection by IHC on tissue sections from the explanted heart of patient A3 and one unaffected control. In the ATTR-CA case, specific staining for TIMP3 showed strong cytoplasmic staining in nearly all cardiomyocytes, with additional weaker expression in the interstitial areas corresponding to amyloid deposits. In contrast, control tissue displayed a patchy staining pattern, characterized by generally weak but locally stronger cytoplasmic expression in cardiomyocytes and occasional focal staining in the interstitial matrix. Overall, staining intensity was markedly higher in the ATTR case compared to the control, consistent with the observed upregulation of TIMP3 in ATTR relative to control (**Figure 3A-B**).

**Figure 3.**
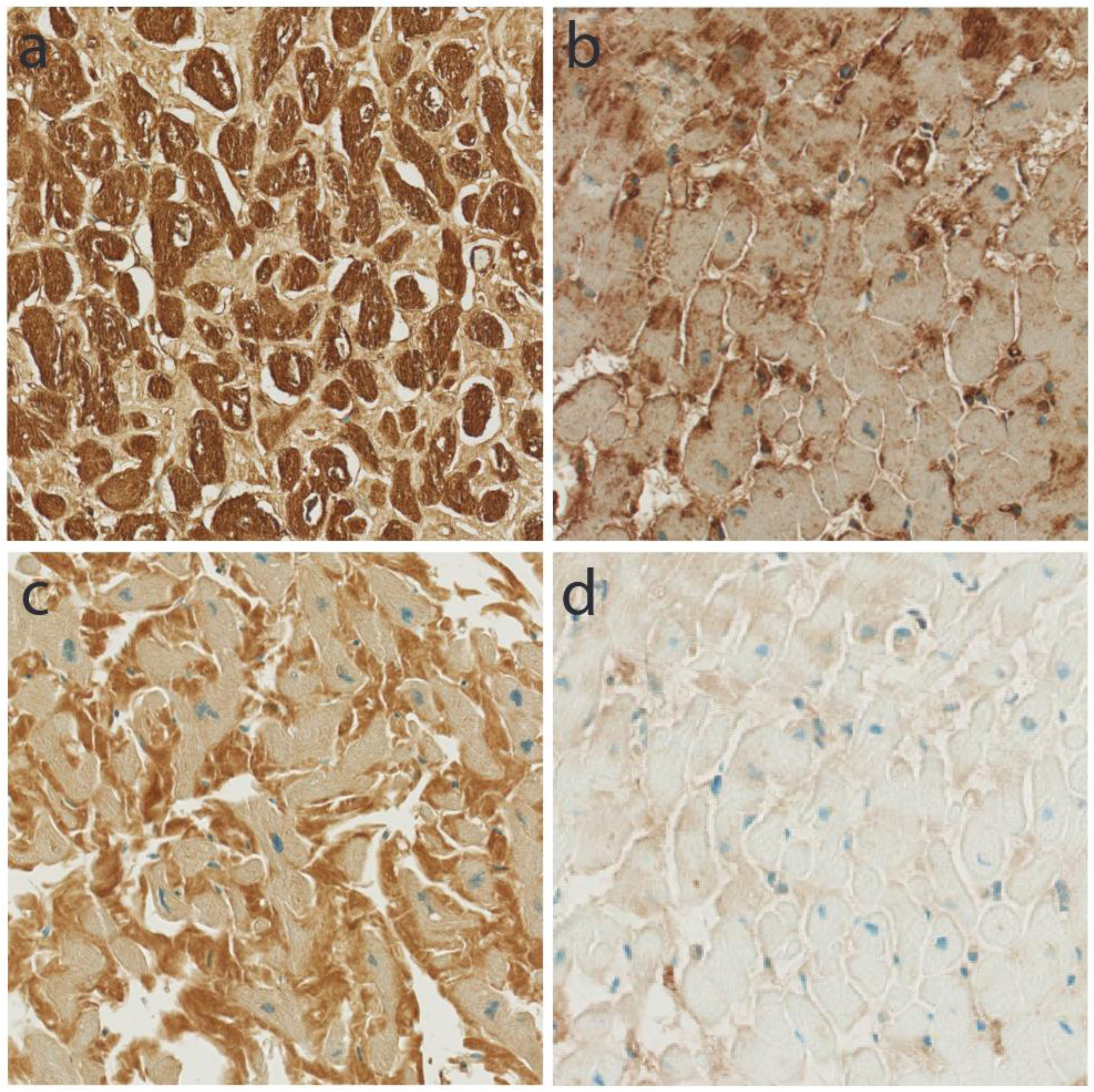
(A–B) TIMP3 immunohistochemistry (IHC) in ATTR and control tissue. TIMP3 shows strong and diffuse cytoplasmic staining in cardiomyocytes and additional weaker expression in amyloid deposits in ATTR, whereas control tissue shows only patchy staining. Original magnification 20x. **(C–D) ADAMTS4 IHC in ATTR and control tissue.** ADAMTS4 shows strong staining in amyloid deposits and weak but uniform cytoplasmic staining in cardiomyocytes in ATTR, with only faint expression in control tissue. Original magnification 20x.

For ADAMTS4, staining in the ATTR case was characterized by a strong and homogeneous signal in the amyloid deposits located between cardiomyocytes. In addition, cardiomyocytes displayed weak but uniform cytoplasmic staining, with comparable intensity across virtually all cells. In contrast, control tissue showed only faint staining, both focally in the interstitial matrix and locally in the cytoplasm of cardiomyocytes, but overall clearly weaker than in the ATTR case. The overall stronger staining of ADAMTS4 in ATTR tissue is consistent with the mass spectrometry data showing ADAMTS4 upregulation in ATTR (**Figure 3C-D**). Extended analyses including AL cases demonstrated that ADAMTS4 showed stronger upregulation in ATTR versus controls compared to AL versus controls, while TIMP3 displayed significant upregulation exclusively in ATTR and not in AL (**Supplementary figure 2, Supplementary Table S7**).

### Heterogeneity in the amyloid-associated proteome

Finally, within the ATTR-CA group, some variability was observed between individual samples. Specifically, three samples (A2, A3 and A6) showed a broader and more intense upregulation pattern compared to the others (**Figure 1C**). This subgroup was marked by increased expression of proteins involved in metabolic and stress-related pathways, including PPAR signaling, lysosome activity, and steroid hormone biosynthesis (**Supplementary Table S2**). This proteomic heterogeneity occurred in addition to the shared proteomic alterations observed across the ATTR-CA cohort. It did not correlate with clinical parameters such as age, gender, genotype, TTR abundance, or amyloid load, suggesting that other unknown disease factors underly this heterogeneity in the amyloid-associated proteome (**Table 1**, **Table 2**).

## DISCUSSION

To investigate the background proteome of cardiac ATTR amyloidosis, we performed quantitative mass spectrometry on whole tissue sections from ATTR-CA cases and unaffected controls. This analysis revealed a distinct proteomic signature characterized by upregulation of ECM remodeling factors, complement components and coagulation proteins.

Looking at ECM remodeling, several MMPs were detected, with significant upregulation of MMP19 and MMP23B, along with TIMP3. This upregulation fits within the hypothesis of matrix remodeling in amyloidosis. Tanaka et al. reported elevated MMP levels in the blood of amyloidosis patients ^25^, and Biolo et al. highlighted their functional relevance in cardiac remodeling and amyloidosis disease progression ^26^. The upregulation of TIMP3 in ATTR amyloidosis plaques in cardiac tissue has previously been noted by Gottwald et al. ^27^ and Kourelis et al. ^28^. We now show that TIMP3 is effectively present in ATTR-CA plaques as well as in cardiomyocytes supposedly producing the protein. Our findings also suggest an involvement of the ADAMTS family of proteins. ADAMTS are zinc-dependent metalloproteinases known for their roles in various diseases, particularly in relation to pathological tissue changes. These enzymes are characterized by their ability to regulate and degrade ECM components, control inflammation and initiate tissue remodeling ^29^.

Immunohistochemistry illustrates this finding in particular for ADAMTS4, and shows markedly increased expression in the ATTR-CA amyloid plaques as compared to unaffected control tissue. Multiple ADAMTSs have previously been implicated in amyloid deposition in the brain ^30,31^. Here, we observed an upregulation of ADAMTS1, ADAMTS4 and ADAMTS5 in ATTR-CA, suggesting that their involvement is not restricted to cerebral amyloid. ADAMTS-like (ADAMTSL) proteins are related to the ADAMTS but lack the protease activity ^32^. Nevertheless, they are known to participate in ECM organization and contribute to the structural integrity of tissues through their role in the formation and maintenance of microfibrils ^33^. ADAMTSL3 showed upregulation in our ATTR cohort.

Taken together, our data indicate that ECM remodeling in ATTR-CA involves not only structural components such as collagen, but also active proteases (ADAMTSs, ADAMTSLs, and MMPs) together with their inhibitor TIMP3. The upregulation of ADAMTS1, ADAMTS4 and ADAMTS5 may reflect a local injury response and associated matrix remodeling. TIMP3, which binds the ECM and regulates protease activity, could serve to restrict excessive matrix degradation ^34^. Protease-inhibitor dynamics seem to contribute to ECM remodeling in amyloidosis, potentially playing an active role in driving tissue dysfunction in ATTR-CA beyond the passive effects of amyloid deposition alone.

Collagen fibers have been shown to envelop amyloid fibrils, contributing to the structural stability of amyloid plaques. A study by Jackson et al. showed collagen to be closely associated with amyloid, protecting fibrils from degradation by macrophages ^35^. Ricagno et al. were able to demonstrate through the use of cryogenic electron tomography (cryo-ET), immune-electron microscopy (IEM) and MS that collagen VI is wrapped around amyloid fibrils in a helical conformation ^36^. Netzel et al. also reported increased levels of collagens in cardiac amyloid proteomes, which were associated with worse clinical outcomes ^17^. These observations are consistent with our proteomic findings indicating an increased presence of multiple collagen types and involvement of procollagen-processing factors in ATTR-CA.

In line with these previous studies, we also observed upregulation of complement components in ATTR-CA. Notably, we found upregulation of C1qA, C1qB and C1qC, C1R, C2, C4A and C4B, C6, C7, C8A, C8B and C8G, C9, CFD, CFH and CFHR1, -2, -3 and -5. Because we used tissue from immunosuppressed patients (heart transplant patients) as controls, we should be careful not to overinterpret the levels of upregulation. However, Lux et al. identified C9 in amyloid deposits in 99.2% of cases, suggesting a critical role for C9 in facilitating the formation of the membrane attack complex (MAC) ^37^. In another publication, he involvement of complement proteins C3, C4A, C5, C9 and CFHR1 and -5 is reported in amyloidosis ^27^. Our findings are consistent with a recent study by Netzel et al., who found complement pathway enrichment in whole tissue proteomic analyses of cardiac ATTR and AL, implicating complement activation in tissue damage and poor clinical outcomes ^17^. Earlier work by Kourelis et al. also provided proteomic evidence for complement involvement in amyloidosis, although these analyses were based on LCM-derived tissue ^28^.

Associated with complement proteins, we found a marked increase in coagulation factors. In literature, not much is known about specific coagulation factors associated with ATTR-CA. In this study, several components of the coagulation pathway were upregulated in the ATTR cohort; F8, F10, F11, F12 and KLKB1 showed marked increases and thrombomodulin, a negative regulator of the coagulation pathway when formed into a complex with protein C and thrombin, was downregulated. Earlier work by Napolitano et al. pointed out coagulation abnormalities in patients with ATTR amyloidosis, who exhibited an increased thrombotic risk that might be linked to amyloid in blood vessels and vascular damage ^38^. The precise implications of the upregulation of coagulation-related proteins in ATTR-CA remain unclear, but they may accompany complement activation. Pathway enrichment analysis supported this observation. While enrichment analyses are not direct proof of functional co-activation, the clustering of complement and coagulation factors within a single pathway is biologically plausible and consistent with the concept of complement-coagulation cross-talk described in other thrombo-inflammatory conditions ^39,40^.

Ultimately, while whole tissue sections prove valuable for studying the amyloid-associated proteome, we observed variability within the ATTR cohort itself. Some samples exhibited significant upregulation of metabolic and stress-related pathways, showing heterogeneity between cases. There was no evident correlation between this upregulation and clinical factors (**Table 1, Supplementary Table 1**). It might be possible that cases with upregulation of these pathways represent more active stages of the disease, with amyloid deposits present in a more dynamic state, eliciting a stronger response. Conversely, samples that did not show upregulation in these pathways may correspond to later stages of the disease. Further investigation would be necessary to confirm some sort of association between proteomic profiles and disease progression. Nevertheless, despite inter-individual variability, our data consistently demonstrate robust upregulation of the pathways discussed above, indicating that these processes represent central and reproducible features of cardiac ATTR amyloidosis.

While amyloid-rich material from LCM deposits is the optimal choice for identifying the amyloid precursor protein in diagnostics, LC-MS/MS of whole tissue sections with the inclusion of adjacent material may contribute to a better understanding of amyloidosis pathogenesis ^16,17^. Moreover, our approach enabled correct classification of all diseased samples as ATTR amyloidosis, indicating that whole tissue sections may offer sufficient diagnostic sensitivity in certain settings, reducing the need for labor-intensive LCM. In tissues with limited amyloid deposits or small biopsies, microdissection of exclusively pathologic tissue is not always feasible. Proteomics on whole biopsies could be a good alternative in these settings. Interpretation however should take into account the presence of adjacent tissue ‘contaminating’ the sample. As such, knowledge of the background proteome is of interest for the pathologist reporting the mass spectrometry results.

Finally, our study has limitations. The small sample size warrants confirmation in larger cohorts where potential heterogeneity can be better interpreted. Additionally, while we used whole tissue sections to capture a comprehensive view of the amyloid-associated proteome, this approach may introduce variability due to differences in tissue composition and amyloid load. While we included controls without histological abnormalities and AL amyloidosis controls, cardiac fibrosis of non-amyloidogenic cause should be included in future work as a non-immunosuppressed control group.

In conclusion, proteomic analysis of whole tissue sections from ATTR amyloidosis revealed robust upregulation of ECM-related proteins, including MMPs, ADAMTSs, and their inhibitor TIMP3, together with complement and coagulation factors. Immunohistochemistry confirmed increased expression of ADAMTS4 and TIMP3 within amyloid plaques. These findings underscore that ECM remodeling and protease-inhibitor dynamics represent reproducible features of cardiac ATTR amyloidosis, and suggest potential avenues for biomarker development and therapeutic intervention.

## Acknowledgements

The authors gratefully acknowledge Irem Kaya for her valuable contribution to tissue processing and preparation, and Malaïka van der Linden for her early coordination and organizational support within the project.

## Author contributions

AV performed manuscript drafting. AV and TMM performed data curation and formal analysis. DVH, SDu and AS contributed to data acquisition and analysis. JS, FR, RG and JVD provided critical review of the manuscript. SD, FI and AD supervised the study and provided overall guidance. All authors have read and approved the final version.

## Ethics statement and patient consent

This study was approved by the Ghent University Hospital’s Ethics Committee (EC: ONZ2022-0301).

