## Supplementary Figures for "Proteomic profiling of whole tissue sections in cardiac ATTR amyloidosis reveals increased extracellular matrix remodeling"

#### SUPPLEMENTARY FIGURE LEGENDS

**Supplementary Figure 1. Extended proteomic analysis including AL cases. (A) Heatmap of all significantly differentially expressed proteins across control, ATTR and AL samples.** Proteins are clustered by unsupervised hierarchical clustering. Control cases are shown on the left, ATTR cases in the middle and AL cases on the right. **(B-D) Heatmaps of selected proteins.** Proteins of interest include those involved in complement activation (B), coagulation cascade (C) and extracellular matrix remodeling (D). Controls are shown on the left, ATTR cases in the middle and AL cases on the right. Upregulation of these pathways is most pronounced in ATTR compared to AL and unaffected controls.

**Supplementary Figure 2. Differential expression of TIMP3 and ADAMTS4 in ATTR and AL.** Volcano plots showing differential expression of TIMP3 and ADAMTS4 in ATTR versus unaffected controls (top), and in AL versus unaffected controls (bottom). The x-axis represents $\log_2$-transformed fold changes ($\log_2$FC) and the y-axis represents $-\log_{10}$ adjusted P-values. Horizontal lines indicate significance tresholds (adjusted $P < 0.05$ and $< 0.01$). ADAMTS4 and TIMP3 are highlighted. ADAMTS4 shows stronger upregulation in ATTR compared to AL, while TIMP3 is significantly upregulated exclusively in ATTR.

### SUPPLEMENTARY FIGURE 1

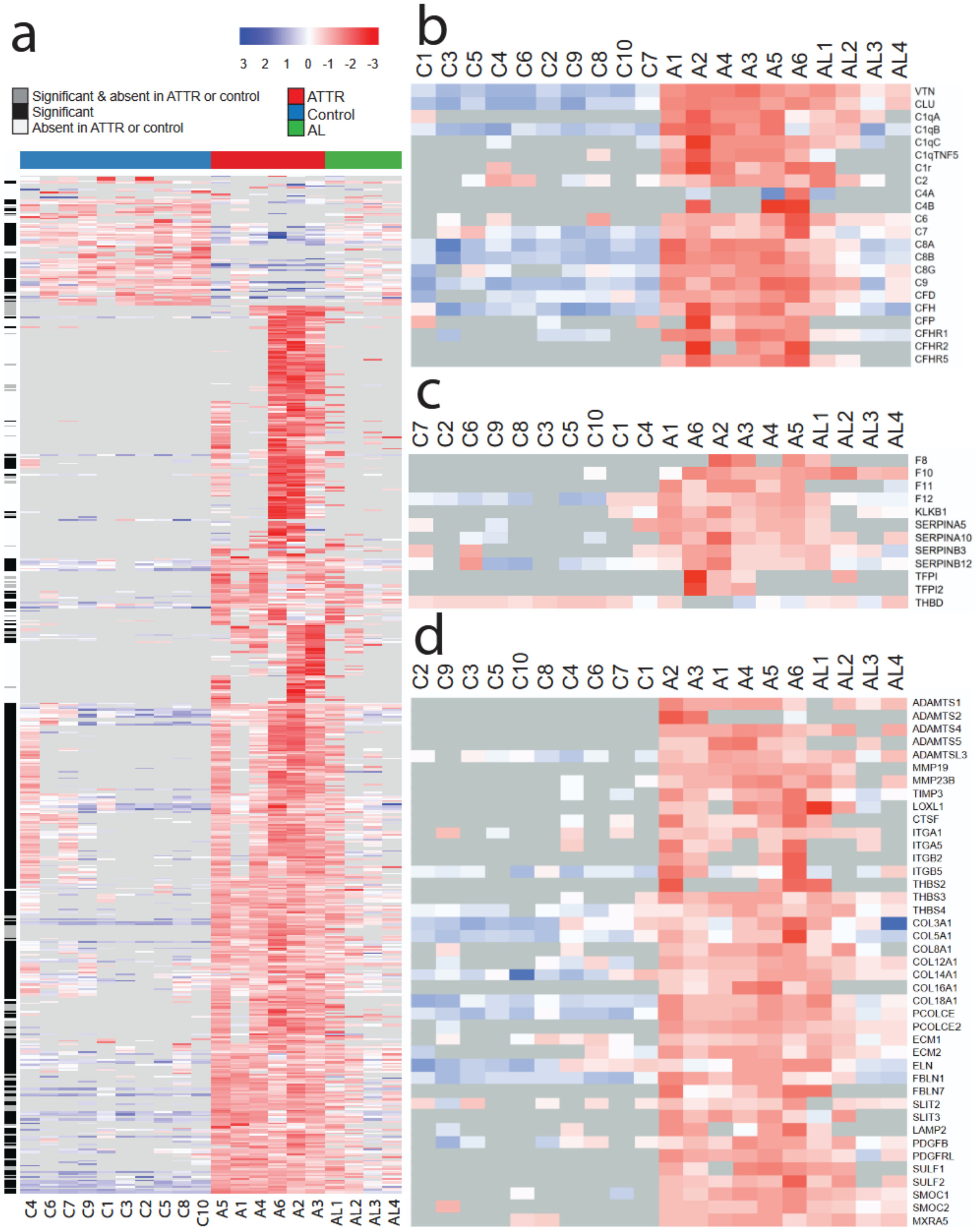

**SUPPLEMENTARY FIGURE 2**

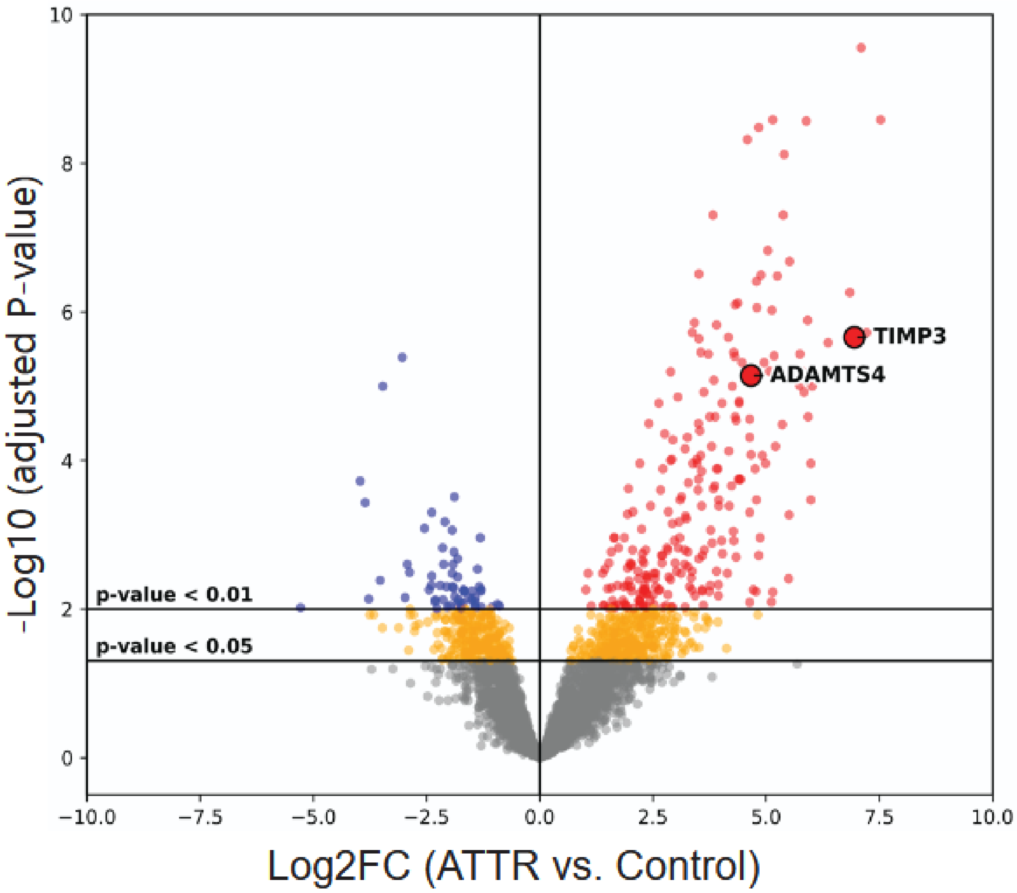


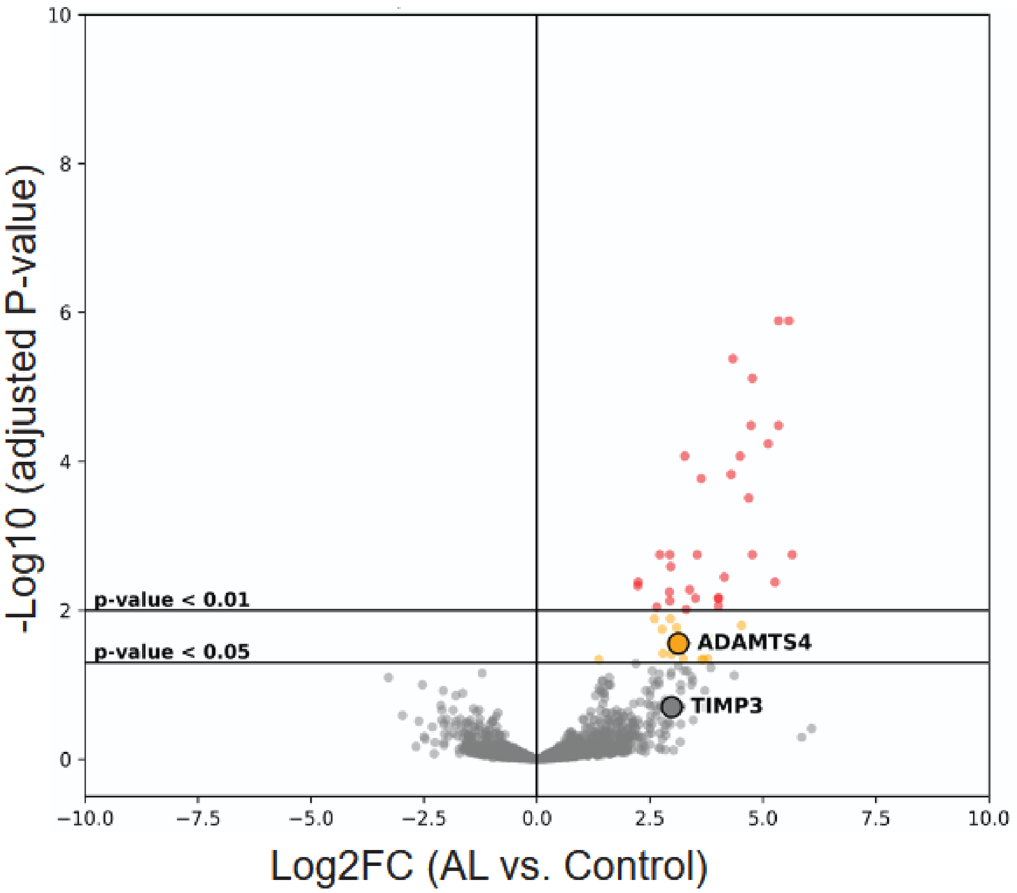
